# Maternal control of RNA decay safeguards embryo development

**DOI:** 10.1101/2025.05.28.656625

**Authors:** Gerardo Del Toro-De León, Maria Sofia Trenti, Varsha Vasudevan, Ursula Krause, Claudia Köhler

## Abstract

As in mammals, the plant embryo is surrounded by maternal tissues that provide protection from the external environment. In angiosperms, a double fertilization process results in the formation of a diploid embryo and a triploid endosperm, both of which develop within maternal sporophytic integuments. Thus, the seed of angiosperms is a combined structure of three genetically distinct components: embryo, endosperm, and maternal integuments. It has long been assumed that the maternal plant influences embryo development, but there is limited molecular evidence for a direct pathway through which the maternal sporophyte affects embryogenesis. Here we show that secondary small interfering RNAs (siRNAs) generated upon exosome impairment lead to embryo abortion through a maternal sporophytic effect. Depletion of the core subunit of the RNA-processing exosome RRP45B (CER7) causes globular embryo arrest connected to ectopic post-transcriptional gene silencing (PTGS). Seed coat expression of *CER7* suppresses seed abortion, demonstrating a maternal sporophytic control of embryo development through an RNA decay safeguard pathway. Our data support a model in which a primary siRNA trigger originates in the maternal integuments of the *cer7* seed coat and drives PTGS amplification in reproductive tissues after fertilization, ultimately leading to seed abortion. In addition, our genetic and molecular data suggest that overloading of AGO1 with siRNAs impairs miRNA function, likely leading to embryo arrest. Our data highlight the complex interplay between maternal and embryonic gene regulation, reinforcing the importance of controlled RNA decay in plant development.

## Introduction

In plants, as in mammals, embryos develop within specialized maternal tissues that provide protection and facilitate nutrient transfer. From basal bryophytes to angiosperms, plants have evolved structures of diverse complexity that support embryo development potentially exerting varying degrees of control over embryogenesis^1^. In angiosperms, the embryo develops surrounded by the endosperm and the seed coat, three genetically distinct components that together constitute the seed. The endosperm is typically a triploid tissue resulting from a second fertilization event, an adaptative reproductive strategy unique to angiosperms. The seed coat is a multilayered envelope of maternal sporophytic origin, derived from the ovule integuments. The complex composition of the seed in angiosperms strongly presumes that both the endosperm and the seed coat influence embryo development.

In the angiosperm model plant *Arabidopsis thaliana*, several lines of evidence indicate that the endosperm is key for embryogenesis, as defects in its development result in embryo arrest and seed abortion^2–4^. For example, embryo patterning requires the peptide molecule EMBRYO SURROUNDING FACTOR 1 (ESF1), which is maternally derived from the central cell, the female gametic precursor cell of the endosperm^5^. Transition from the coenocytic to the cellular stage of endosperm development is crucial for embryo survival and implicated to establish desiccation tolerance in the embryo^2^. Furthermore, regulated endosperm breakdown by the endosperm-specific *ZHOUPI* gene is important for embryonic epidermal development^6^.

In contrast, little is known about how the maternal sporophytic tissues impact embryogenesis. Besides being a physical protective structure, little evidence implicates the maternal integuments in a more direct role during embryo development. Recent reports suggest that the seed coat could potentially work as a communication barrier regulating the movement of molecules across seed compartments. For example, auxin is actively transported from the integuments to the suspensor^7^, the ephemeral basal cell of the embryo and the only known symplastic connection to the surrounding maternal tissues. Other molecules have been suggested to be transferred from the seed coat to the fertilization products, but the evidence is scarce or indirect^8–10^. In genetics, the influence of the integuments on embryogenesis implies a sporophytic maternal genetic effect. Sporophytic maternal mutations with embryo defects reveal the function of the impaired gene in maternal tissues relevant to embryogenesis, which translates into a non-Mendelian penetrance of the mutant phenotypes. Only a handful of reports have provided genetic evidence supporting sporophytic maternal control over embryo development^11–14^. However, the mechanisms are poorly understood. Here we report the discovery of a sporophytic maternal pathway surveilling embryo development through an RNA processing decay mechanism. Our genetic and molecular data demonstrate that mutations in *ECERIFERUM7* (*CER7*)/*RRP45B*, a core component of the RNA exosome, are embryo lethal with a sporophytic maternal effect. The exosome complex is an evolutionarily conserved holoenzyme in eukaryotes responsible for 3′-5′ RNA decay. Nuclear and cytoplasmic forms of the exosome share a core complex constituted by a barrel-shaped structure of nine subunits associated to the ribonucleolytic subunit DIS3/RRP44^15,16^. Mutations in the core components of the exosome are often lethal in *Arabidopsis thaliana*. Disruption of the RRP41L subunit delays germination and affects early seedling development^17^, while mutations in *RRP41*, *RRP42*, *DIS3*/*RRP44A* and *RRP4* have been reported to cause female gametophyte and embryo defects^18–21^. Interestingly, mutations in the core exosome subunit encoding gene *CER7* result in severe seed inviability^22,23^, but the underlying cause of this defect remains unknown. The *cer7* mutant was initially identified by its wax-deficient outer cuticular surfaces, which result in glossier or brighter stem phenotypes^24^. This phenotype is caused by the accumulation of secondary small interfering RNAs (siRNAs) targeting the key wax biosynthetic gene *CER3*, causing reduced *CER3* RNA levels^22,25–27^. Disruption of RNA metabolic processes often results in aberrant transcripts entering the post-transcriptional gene silencing (PTGS) pathway, causing reduced transcript levels of endogenous genes^28–30^.

We found that the exosome prevents the accumulation of aberrant transcripts in the seed coat, capable of triggering RNA silencing in the fertilization products impairing embryo development. Our data suggest that in the *cer7* mutant, RNA molecules derived from sporophytic maternal tissues induce embryo arrest through ectopic RNA silencing, implying RNA transport from the maternal sporophyte to the embryo.

## Results

### *cer7* mutants arrest at the globular stage of embryo development

To characterize the reduced seed viability observed in *CER7/RRP45B* mutants, we analyzed two independent loss-of-function alleles, *cer7-3* (SAIL_747_B08) and *cer7-4* (GK_089C02) (**Supplementary Fig. 1**)^22,23^. Homozygous mutants of both *cer7* alleles produced strongly wrinkled seeds (**Supplementary Fig. 2A**). To accurately determine seed abortion rates, we performed manual crosses and quantified seed abortion at 14 days after pollination (DAP). We found that over 90% of developing seeds displayed a size comparable to wild-type (WT) siblings within the same silique but lacked green pigmentation, suggesting embryonic arrest; we refer to these as ‘aborted seeds’. (**Supplementary Fig. 2**). To further characterize defects associated with seed abortion, we assessed embryo development at 2, 3, 5, and 7 DAP. At 2 DAP most developing seeds were at the 1-2 cell embryo stage in WT and both *cer7* mutant alleles. At 3 DAP ∼25% of WT embryos were at the preglobular and 75% at the globular stage, while >85% of the *cer7* embryos of both alleles remained at the preglobular stage (**Fig. 1A-B**), indicating a slight delay in development compared to WT. At 5 and 7 DAP, over 95% of wild-type embryos had reached the heart and torpedo stage, respectively, whereas embryos from both *cer7* mutants remained at the globular stage with no apparent cell division defects. These findings show that *cer7* mutant seeds exhibit a strong developmental arrest at the early to mid-globular stages of embryogenesis (**Fig. 1A-B**). We did not detect defects in endosperm development in both *cer7* alleles, proliferation and cellularization occurred at same speed as in WT, indicating that defects in *cer7* are primarily restricted to embryo development (**Fig. 1C**). We therefore concluded that *cer7* shows seed inviability mainly due to a globular embryo arrest.

**Fig. 1:**
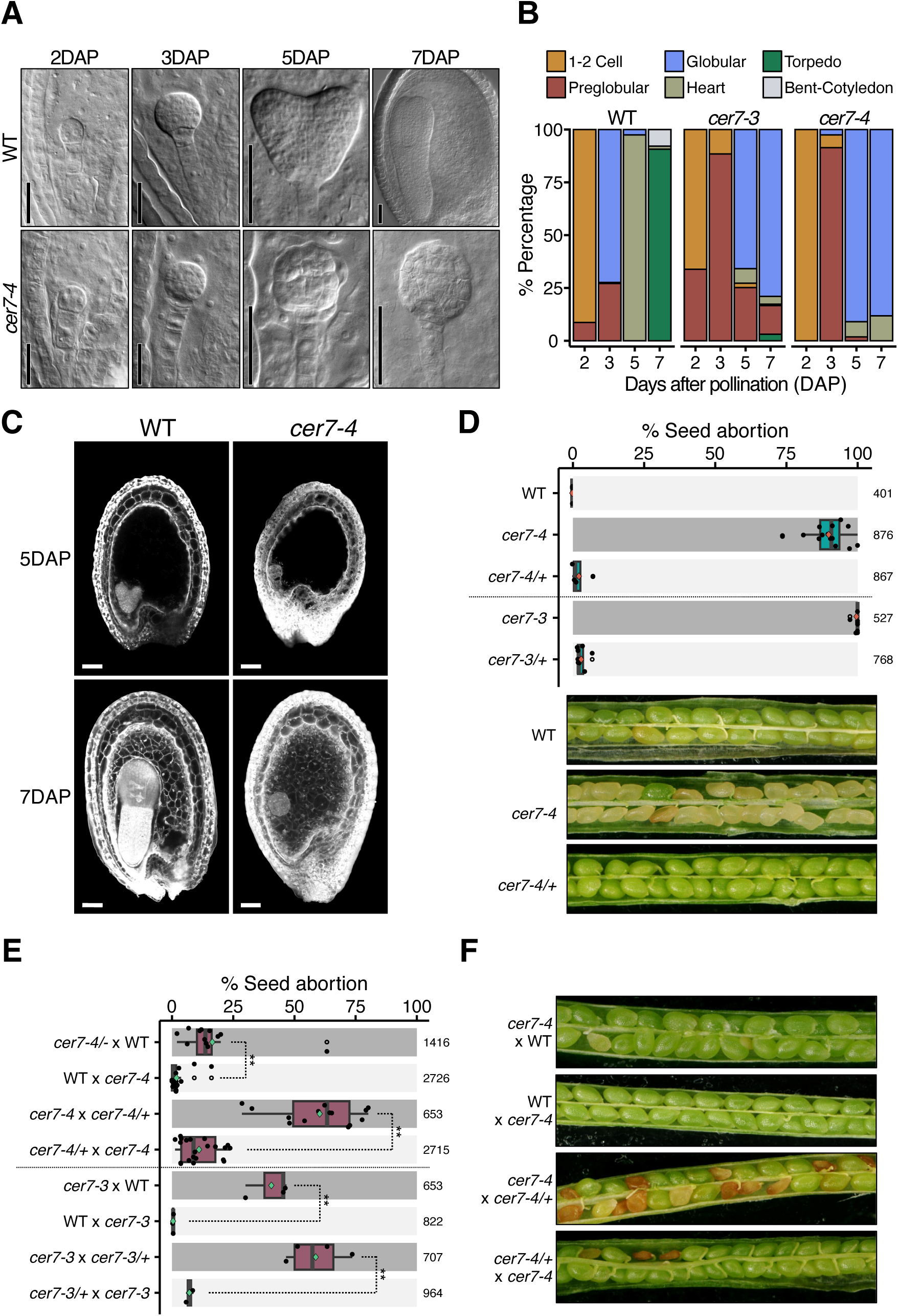
Phenotypic and genetic characterization of *cer7* mutants. (**A**) Nomarsky images of embryos in whole mount cleared seeds of wild-type (WT) Col-0 and *cer7-4* at 2, 3, 5 and 7 days after pollination (DAP). (**B**) Quantification of embryo stages in WT and *cer7* alleles. (**C**) Endosperm cellularization in WT and *cer7-4* determined by Feulgen staining at 5 and 7 DAP. (**D**) Quantification of seed abortion in hand self-pollinated heterozygous plants at 14 DAP (top panel), representative images of siliques (bottom panel). (**E**) Quantification of seed abortion at 14 DAP in indicated genetic crosses to determine the maternal effect in *cer7* mutants. (**F**) Representative images of siliques quantified in (E). Bar color code for D and E: dark gray denotes sporophytic maternal plant homozygous for *cer7*; light gray denotes WT and heterozygous *cer7* sporophytic maternal plant. Pairwise t-test, **P-value < 0.01; * P-value < 0.05; ns, not significant. Scale bar 50 µm (A and B).

### *cer7* mutants show a sporophytic maternal eOect on embryo lethality

To determine whether the seed phenotype of *cer7* follows a recessive Mendelian segregation, we first quantified seed abortion in heterozygous plants. Surprisingly, heterozygous plants did not exhibit seed abortion in either of the two independent alleles tested (**Fig. 1D**). Segregation analysis of the progeny showed the expected 1:2:1 genotypic ratio of a classic recessive mutation (16*+/+*:49*+/-*:16*-/-*, *P*-value >0.05, Chi-square test), arguing against gametophytic defects biasing phenotypic proportions. We therefore hypothesized that *cer7* embryo arrest is caused by a sporophytic maternal effect. To test this hypothesis, we reciprocally crossed homozygous *cer7* mutants with WT or pollen from heterozygous mutants and scored abortion at 14 DAP. We found that *cer7* homozygous mutants crossed with WT pollen resulted in 17% and 40% of seed abortion for *cer7-4* and *cer7-3*, respectively. In contrast, WT maternal plants crossed with *cer7* pollen showed normal seed viability (**Fig. 1E-F**). Interestingly, when pollinating homozygous *cer7* plants with heterozygous pollen we observed 60% of aborted seeds for both alleles tested, while the genetically identical progeny derived from crossing heterozygous plants with homozygous pollen showed 11% and 7% of aborted seeds for *cer7-4* and *cer7-3*, respectively (**Fig. 1E-F**). Collectively, these data reveal that homozygous and heterozygous *cer7* embryos abort at high rates only when developing within homozygous maternal *cer7* mutants, demonstrating that *cer7* is a sporophytic maternal effect mutant with embryo defects. Importantly, our genetic analysis showed 60% of seed abortion when homozygous maternal *cer7* mutants were pollinated with pollen from heterozygous mutants, compared to >90% of seed abortion from self-pollinated homozygous plants, indicating an additional zygotic component on seed abortion. Since both *cer7* alleles consistently exhibited equivalent phenotypes, subsequent experiments were conducted using *cer7-4*.

### *cer7* embryo lethality is triggered by maternally controlled PTGS

Disruption of *CER7* has been implicated to cause aberrant transcripts entering the PTGS pathway, causing reduced transcript levels of endogenous genes^22,25–27^. We therefore hypothesized that seed abortion in *cer7* could also be explained by ectopic PTGS. To test this idea, we introduced *cer7* into *sgs3* and *rdr6* mutant backgrounds, two key players in RNA silencing^31^. We indeed found that *cer7;sgs3* and *cer7;rdr6* double mutants fully rescued seed abortion (**Fig. 2A-B**), demonstrating that embryo arrest in *cer7* is dependent on aberrant PTGS.

**Fig. 2:**
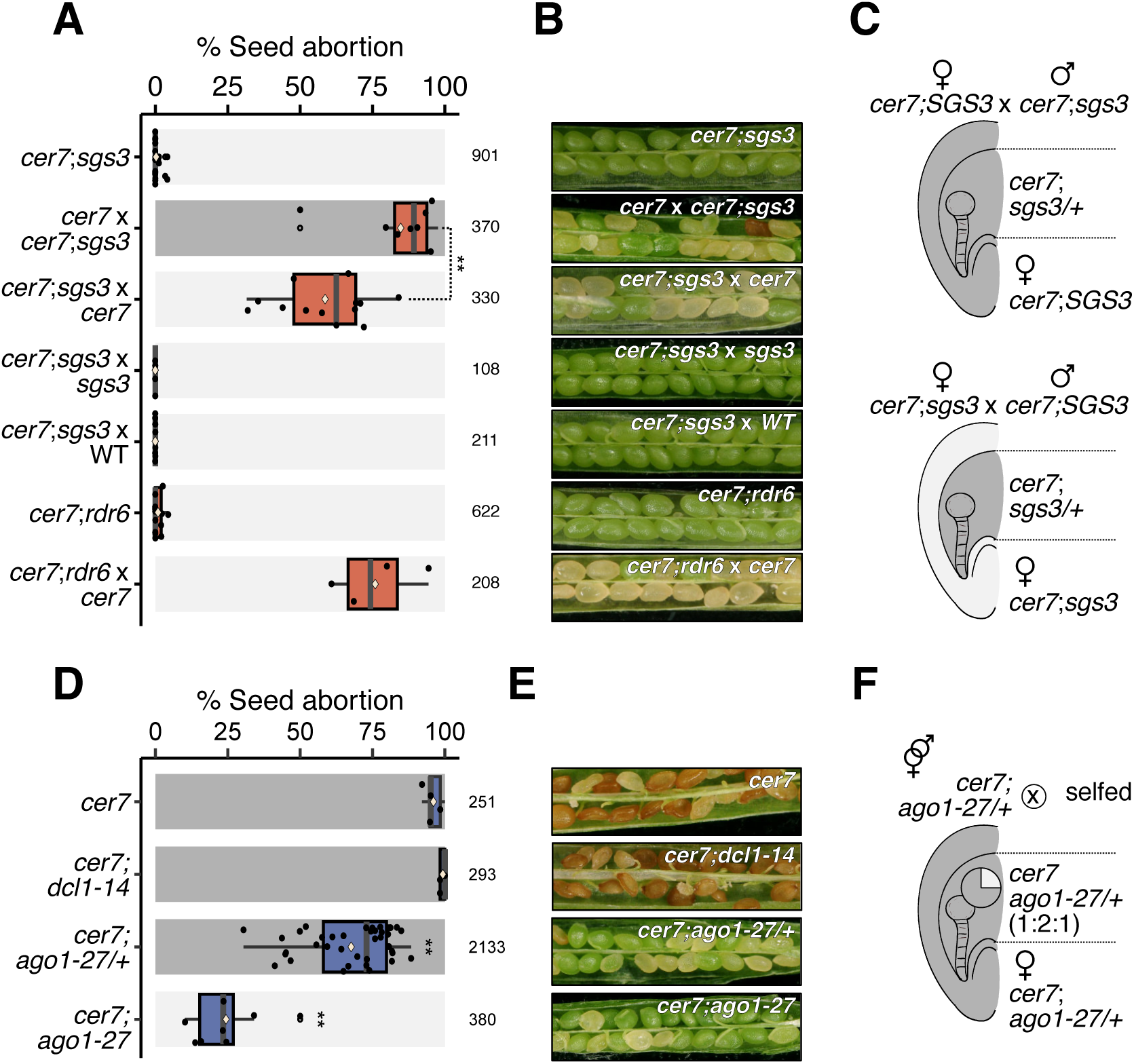
Seed abortion in *cer7* is dependent on maternally controlled PTGS. (**A**) Quantification of seed abortion at 14 days after pollination (DAP) in the indicated genetic crosses. Numbers of analyzed seeds are indicated on the right side of the graph. (**B**) Representative images of siliques quantified in (A). (**C**) Scheme depicting the expected PTGS amplification in reciprocal crosses. Dark grey indicates active PTGS amplification; light grey indicates impaired amplification. Top panel: *cer7;SGS3* sporophytic maternal plant and progeny amplify PTGS. Bottom panel: *cer7;sgs3* maternal plant with PTGS impairment; progeny retains amplification. (**D**) Quantification of seed abortion at 14 DAP in the indicated genetic crosses. Numbers of analyzed seeds are indicated on the right side of the graph. (**E**) Representative images of siliques quantified in (C). Bar color code for B and E: Dark grey indicates PTGS amplification by the *cer7* homozygous sporophytic maternal plant; light grey indicates impaired PTGS amplification in the sporophytic maternal plant. (**F**) As in (A), scheme depicting the expected PTGS amplification in selfed *cer7;ago1-27/+* plants. The sporophytic maternal plant and the majority of the progeny amplify PTGS (dark grey). A subset of progeny (*ago1-27* homozygotes, ∼25%) is expected to show impaired PTGS amplification, indicated in light grey in the accompanying pie chart. Pairwise t-test, **P-value < 0.01; * P-value < 0.05; ns, not significant.

Amplification of siRNAs requires the synthesis of double-stranded RNAs (dsRNAs) mediated by SGS3-RDR6, which are later processed by DCL4/2 to produce 21/22-nt siRNAs^31,32^. Since we determined that the *cer7* seed phenotype is established by a sporophytic maternal effect, we aimed to analyze whether PTGS-mediated seed collapse was likewise maternally controlled. To test this idea, we reciprocally crossed homozygous *cer7* mutants with *cer7;sgs3 and cer7;rdr6* double mutants to deplete PTGS amplification in either the sporophytic maternal or zygotic tissues and scored seed abortion at 14 DAP. Embryos resulting from crosses between maternal *cer7* plants and *cer7;sgs3* pollen develop within a sporophyte capable of amplifying siRNAs, as SGS3 is functional in the maternal sporophytic tissues (**Fig. 2C**). Accordingly, we observed an average of 85% of aborted seeds in this cross. In contrast, embryos resulting from the reciprocal cross develop within a *cer7;sgs3* maternal sporophyte unable of amplifying PTGS, but the fertilization products retain this ability given the functional *SGS3* copy inherited from the paternal parent (**Fig. 2C**). This cross resulted in 59% aborted seeds, which was significantly less than in the reciprocal cross. Similar results were obtained with *cer7 and cer7;rdr6* reciprocal crosses (**Fig. 2A-B**). These observations indicate that *cer7;sgs3/rdr6* maternal sporophytes are capable of inducing PTGS-mediated embryo arrest, meaning that siRNA amplification can also occur in the fertilization products. These results suggest that the trigger for PTGS in the maternal sporophytes is upstream of SGS3-RDR6, likely an aberrant RNA resulting from deficient exosome function. Our observations imply that the trigger that induces aberrant PTGS is generated in the sporophytic maternal tissues in alignment with the sporophytic maternal-effect, and is capable to reach the embryo for subsequent RNA silencing amplification, leading to *cer7* embryo arrest. In agreement with this notion, we observed higher seed abortion rates in *cer7* x *cer7;sgs3* compared to *cer7;sgs3* x *cer7* (85% *versus* 59%), suggesting that PTGS amplification in the *cer7* maternal sporophyte contributes to an increased dosage of ectopic siRNAs resulting in higher seed abortion frequency. We therefore concluded that *cer7* seed abortion is the result of PTGS, which is maternally triggered in sporophytic tissues and amplified in both the maternal sporophyte and the fertilization products likely through RNA movement across seed compartments.

### AGO1 is required for PTGS-mediated *cer7* embryo arrest

AGO1 is the main effector of PTGS. We therefore asked whether *cer7* PTGS-dependent seed abortion was mediated by AGO1. To test this idea we introduced *cer7* into the *ago1-27* mutant, a weak allele specifically impaired in sense PTGS (S-PTGS)^33^. Indeed, the *cer7*;*ago1-27* double mutant fully suppressed seed abortion, implicating RNA silencing through AGO1 (**Fig. 2D-E**). We then assessed whether *ago1-27* is required in the sporophyte or in the fertilization products to rescue *cer7* seed abortion. To test this, we scored seed abortion in *cer7*;*ago1-27/+* heterozygous plants. We anticipated that if *ago1-27* suppresses seed abortion zygotically, at least 25% of the progeny should be viable (**Fig. 2F**). Indeed, we found that self-pollinated *cer7*;*ago1-27/+* plants showed an average of 32 % viable seeds, in comparison to 2% in *cer7* homozygous plants (**Fig. 2D-E**). These data indicate that AGO1 acts in the fertilization products to induce *cer7* seed abortion.

Finally, we explored whether aberrant PTGS in *cer7* was triggered by microRNA activity. Little is known about the mechanisms by which PTGS is initiated in endogenous genes. However, similar to *TAS* genes, a subset of endogenous transcripts prone to PTGS amplification are targets of microRNAs^34,35^. To test this idea, we introduced *cer7* into the *dcl1-14* background, a viable hypomorphic allele of DCL1 which encodes a key RNase III enzyme required for miRNA biogenesis. The *cer7;dcl1-14* double mutant showed equivalent abortion levels to the *cer7* single mutant, arguing against microRNA-triggered PTGS (**Fig. 2D-E**). Nevertheless, *dcl1-14* mutant is a weak allele and retains low levels of microRNA activity^36^; therefore, we cannot exclude the possibility that leaky microRNA activity in *dcl1-14* is sufficient to induce PTGS amplification in *cer7*.

### *CER7* activity in the seed coat rescues seed abortion

Our data strongly suggest that an aberrant transcript or a subset of transcripts accumulate in *cer7* sporophytic maternal tissues and reach the embryo, leading to PTGS amplification and seed abortion. We reasoned that by expressing *CER7* in the seed coat we could prevent the generation and movement of aberrant transcripts entering PTGS, suppressing embryo arrest. To test this hypothesis, we generated complementation lines expressing *CER7-GFP* under the control of promoters that are specifically active in the seed coat (**Fig. 3A-E**). Constitutive expression of *CER7-GFP* under the *RPS5A* promoter fully suppressed both the glossy stem phenotype and seed abortion in all lines tested, indicating that *CER7-GFP* is functional (**Fig. 3A, E, F**). We then expressed *CER7-GFP* under control of the *BANYULS* (*BAN*) promoter, which is specifically active in the endothelium, the innermost cell layer of the inner integument (ii1)^37,38^. We found that *BAN::CER7-GFP* expression caused a significant reduction of seed abortion in 27% of the lines tested (**Fig. 3B, G**). Since all the three layers of the inner integument are symplastically connected^9^, we reasoned that aberrant transcripts could easily move from the outermost layers and potentially overload endothelium *CER7-GFP* activity. We therefore expressed *CER7* under the *deltaVPE* (*dVPE*) promoter, which is specifically active in the outer two cell layers of the inner integument (ii2 and ii3)^39^. Expression of *dVPE::CER7-GFP* resulted in a significant reduction of seed abortion in 40% of the lines tested (**Fig. 3C, H**). We then analyzed the combined complementation effect of *CER7-GFP* under control of *dVPE* and *BAN* (ii1, ii2, ii3) by crossing those lines that showed little to no rescue. We observed a strong additive effect, as all tested plants expressing both constructs showed a significant reduction of seed abortion (**Fig. 3D, I**). Importantly, neither the *dVPE* nor the *BAN* promoter *CER7-GFP* expression rescued the glossy stem phenotype (**Fig. 3E**). These results demonstrate that the activity of *CER7* is required in the seed coat to prevent formation of aberrant transcripts, consistent with the sporophytic maternal-effect of *cer7*.

**Fig. 3:**
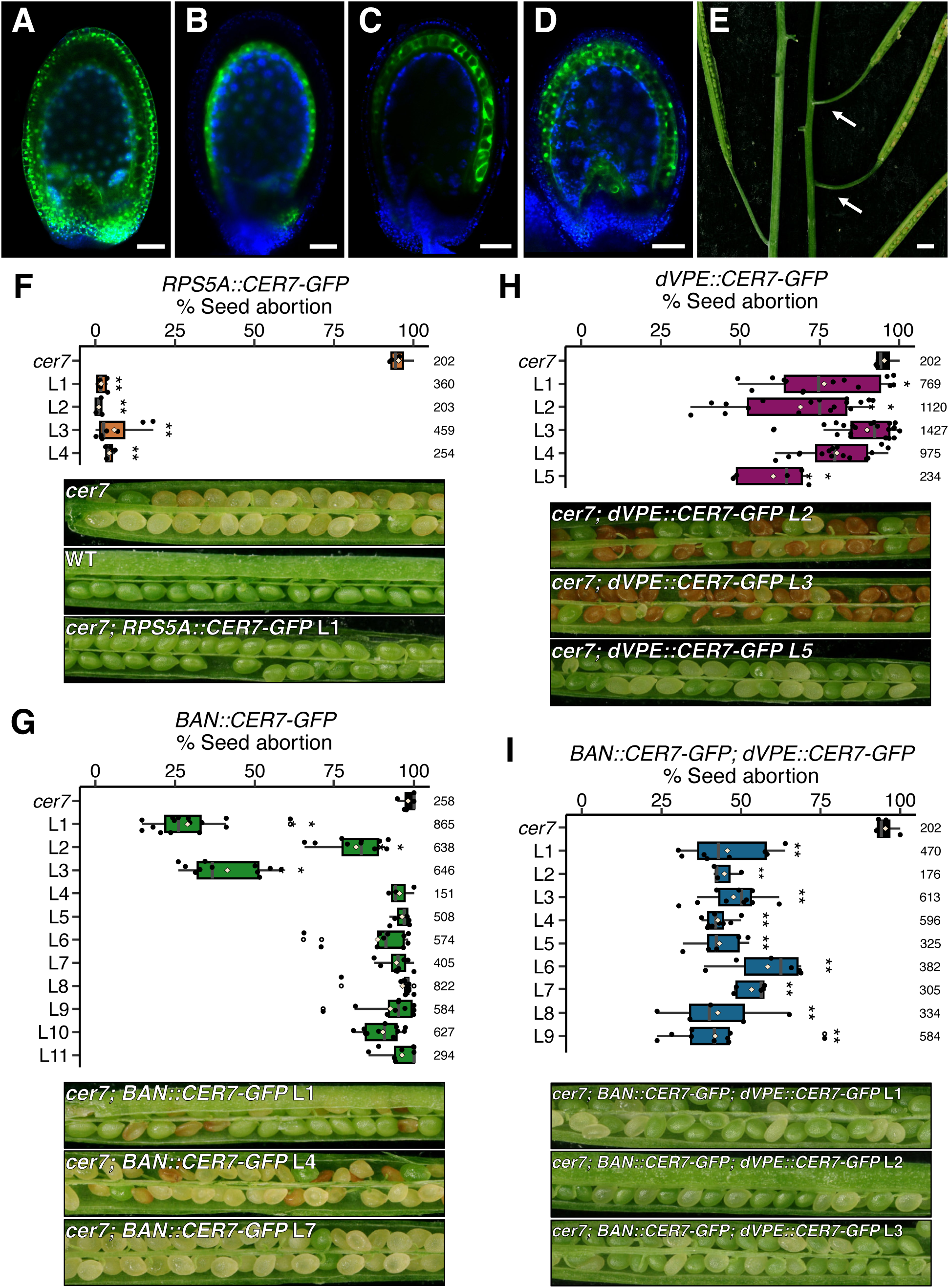
Seed coat expression of *CER7* suppresses *cer7* seed abortion. (**A–D**) *CER7-GFP* expression driven by the constitutive *RPS5A* promoter (A), the seed coat-specific *dVPE* promoter (B), the *BAN* promoter (C), or a combination of both *dVPE* and *BAN* promoters (D). Green represents GFP expression, blue shows autofluorescence. Scale bar 50 µm. (**E**) Stems of 6 week old *cer7;pRPS5A::CER7-GFP* (left) and *cer7;pBAN::CER7-GFP;pdVPE::CER7-GFP* (right). White arrows indicate glossy stem phenotype. Scale bar 200 µm. (**F-I**) Quantification of seed abortion in mature siliques in *cer7* mutants expressing *CER7-GFP* under the constitutive *RPS5A* promoter (F), the seed coat-specific inner integument (ii1) *BAN* promoter (G), the ii2-3 specific *dVPE* promoter (H) and the combined expression of d*VPE* and *BAN* promoters (ii1-3) (I). Representative images of the indicated siliques depicted in the bottom panels. Numbers of analyzed seeds are indicated on the right side of the graph. DAP, days after pollination. L, individual transgenic line. Pairwise t-test, **P-value < 0.01; * P-value < 0.05; ns, not significant.

### PTGS response in *cer7* developing seeds

We have shown that *cer7* embryo lethality is dependent on maternally controlled PTGS, which implies RNA silencing mediated by ectopic accumulation of siRNAs. We therefore hypothesized that a gene or group of genes relevant for embryo development were targeted by siRNAs produced both in sporophytic and fertilization products leading to embryo arrest. We thus aimed to identify downregulated genes targeted by siRNAs in *cer7* seeds. For this purpose, we generated sRNA-seq and RNA-seq libraries in triplicates of WT, *cer7*, *sgs3,* and *cer7;sgs3* seeds at 3 DAP (**Supplementary Fig. 3-4, Supplementary Table 1-2**). First, we asked whether coding genes were indeed ectopically generating siRNAs in *cer7*. Because PTGS is mediated by 21-nt and 22-nt siRNAs, we fragmented the genome in 100bp windows and identified significantly enriched regions with 21/22-nt siRNAs in *cer7* samples (LFC > 0; FDR <0.05). As expected, over 90% of the 21/22-nt sRNA-enriched windows in *cer7* overlapped with coding genes. Since SGS3 is required for PTGS and *sgs3* suppressed the *cer7* mutant phenotype, we then selected those loci that lost siRNAs in the *cer7;sgs3* double mutant (**Supplementary Fig. 5A-B**). We analyzed whether the distribution of 21/22-nt siRNA was present on both strands of coding sequences, a hallmark of PTGS silencing, and identified 19 genes targeted by secondary siRNAs in *cer7* seeds (**Fig. 4A, Supplementary Fig. 5 and Supplementary Dataset 1**). Next, we tested whether ectopic 21/22-nt siRNA accumulation correlated with reduced gene expression. We conducted differential gene expression analysis (LFC ±0.58; FDR <0.05, **Supplementary Fig. 6-7, Supplementary Dataset 2**), but found only three genes that were downregulated and accumulated secondary 21/22-nt siRNAs in *cer7* seeds: *AGO1*, *XYLOGLUCAN ENDOTRANSGLUCOSYLASE 9* (*XTH9*) and *BETA GALACTOSIDASE 18* (*BGLU18*) (**Fig. 4B**). To evaluate the potential relevance for early embryo development, we analyzed gene expression in seed compartments using public datasets^40^. *AGO1* and *XTH9* genes are expressed in embryonic tissues, whereas *BGLU18* showed no detectable expression, making its involvement in embryo arrest in *cer7* mutants unlikely (**Fig. 4C**). *XTH9* encodes an enzyme involved in cell wall remodeling and expansion. However, since *xth9* mutants have no reported seed defects^41^, reduced *XTH9* expression is unlikely to contribute to the *cer7* phenotype. Notably, *ago1* null mutants show strong pleiotropic phenotypes and infertility, demonstrating essential functions in development^42,43^. However, early globular embryo arrest has not been reported in *ago1* mutants to date. Therefore, we concluded that PTGS directed on individual genes alone is not sufficient to explain the early globular embryo arrest in *cer7*, suggesting that addi-onal factors contribute to seed abor-on.

**Fig. 4:**
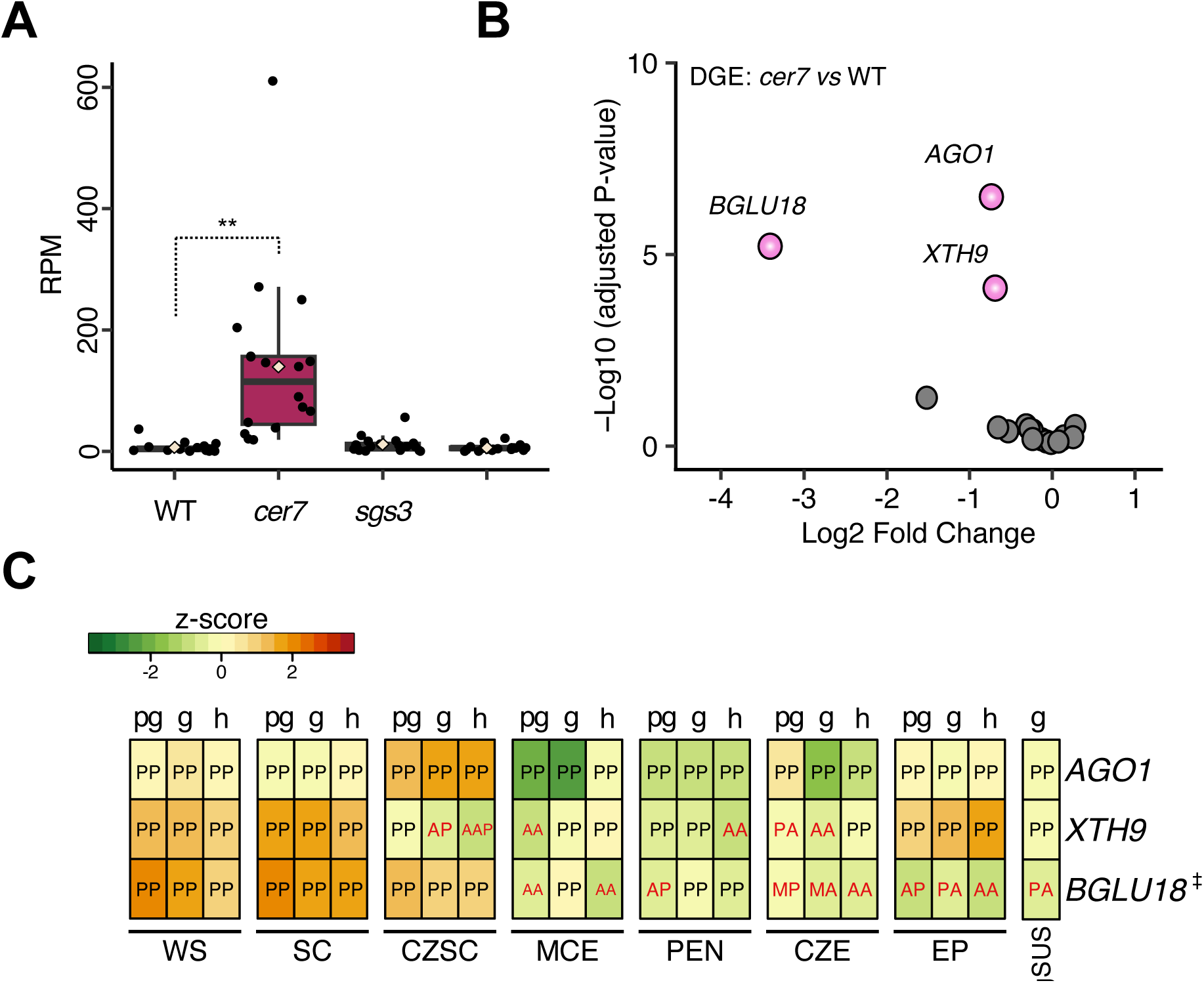
PTGS in *cer7* developing seeds at 3 DAP. (**A**) sRNA read counts (RPM) for 21/22-nt siRNA accumulating genes in *cer7* at 3 DAP seeds of *cer7*. (**B**) RNA-seq of 3 DAP seeds showing differential expression of genes accumulating 21/22-nt siRNAs in *cer7*. Pink color indicates genes with reduced transcript levels and grey indicated genes with no significant change. (**C**) Expression level of 21/22-nt siRNA accumulating genes in different seed compartments along development. ‡ Genes with low or no expression in the embryo. Letters within heatmap boxes indicate MAS5 detection calls: P, present; A, absent; M, marginal. Letters above heatmap boxes indicate developmental stages: pg, preglobular; g, globular; h, heart-stage; lc, linear cotyledon; m, mature green. WS, whole seed; SC, seed coat; CZSC, chalazal seed coat; MCE, micropylar endosperm; PEN, peripheral endosperm; CZE, chalazal endosperm; EP, embryo proper; SUS, suspensor (Belmonte et al. 2013). Pairwise t-test, **P-value < 0.01; * P-value < 0.05; ns, not significant.

### AGO1 loading of a subset of miRNAs is compromised in *cer7*

We showed that PTGS-mediated *cer7* embryo arrest requires AGO1 function (**Fig. 2D-F**) and that *AGO1* transcripts were targeted by siRNAs in *cer7*, resulting in a 50% reduction of transcript levels compared to WT (**Supplementary Fig. 7A**). Previous work revealed that ectopically accumulating 21-nt siRNAs compete with miRNAs for loading into AGO1^44^. Because miRNAs are instrumental for embryo development^45^, we hypothesized that *cer7* mutants are compromised in miRNA function due competing ectopic siRNAs. Supporting this idea, the sporophytic maternal effect observed in *cer7* echoes the phenotype of the *short integuments1* (*sin1*) mutant, an allele of *DCL1* that like *cer7* shows maternal sporophytic embryo arrest^13^. Moreover, loss of *MIR167* function also causes a sporophytic maternal effect with embryonic defects, strikingly similar to those observed in *cer7*^14^. To test whether indeed miRNA loading was compromised in *cer7*, we analyzed the populations of 21-nt siRNAs in our sRNA-seq seed samples and found a significant reduction of miRNAs in *cer7* compared to the other genotypes (**Fig. 5A**). We found a higher proportion of ribosomal RNA-derived 21-nt sRNAs (ri-siRNAs) in *cer7,* in line with previous reports showing increased accumulation of 21-nt ri-siRNAs in mutants impaired in RNA degradation^14^ (**Fig. 5A**). To directly test which populations of sRNAs were loaded into AGO1 in the *cer7* mutant, we performed small RNA immunoprecipitation followed by high-throughput sequencing (RIP-seq) of AGO1 from 2-3 DAP WT and *cer7* siliques in duplicates (**Supplementary Fig. 8, Supplementary Table 3**). We first analyzed the size distribution of our RIP-seq libraries and indeed found an enrichment of 21-nt sRNAs and a depletion of 24-nt sRNA as expected for sRNAs associated with AGO1 (**Fig. 5B**). The great majority of the 21-nt sRNAs corresponded to miRNAs in all samples; however, we observed a higher proportion of 21-nt siRNAs derived from protein coding genes but not ri-siRNAs in *cer7* samples and a proportional reduction of miRNAs (**Fig. 5C**). To examine differences in AGO1 loading of sRNAs, we fragmented the genome into 100bp windows and identified regions significantly enriched or depleted of 21/22-nt sRNA in *cer7* compared to WT (LFC ±0; FDR <0.05). Consistent with our sRNA-seq analysis from seeds, all genes identified as enriched for 21/22-nt sRNAs were also enriched in our AGO1 IP data (**Supplementary Fig. 9, Supplementary Dataset 3**). This observation confirms that a subset of genes generates ectopic siRNAs that compete for AGO1 loading with miRNAs. To analyze whether specific miRNAs were depleted from AGO1 in *cer7*, we analyzed the identified windows significantly depleted of 21/22-nt sRNAs in *cer7* compared to WT. Indeed, we found that these regions correspond to a subset of miRNAs that were significantly less abundant in AGO1-IP samples from *cer7* (**Supplementary Dataset 4**). To confirm whether these miRNAs were recovered in *cer7;sgs3*, we analyzed their relative abundance across our sRNA-seq seed samples and indeed found that *cer7* showed a significant reduction of these miRNAs compared to the other genotypes (**Fig. 5D**). We therefore concluded that in *cer7* ectopic accumulation of 21/22-nt siRNAs derived from PTGS impairs loading of a subset of miRNAs onto AGO1, compromising its function.

**Fig. 5:**
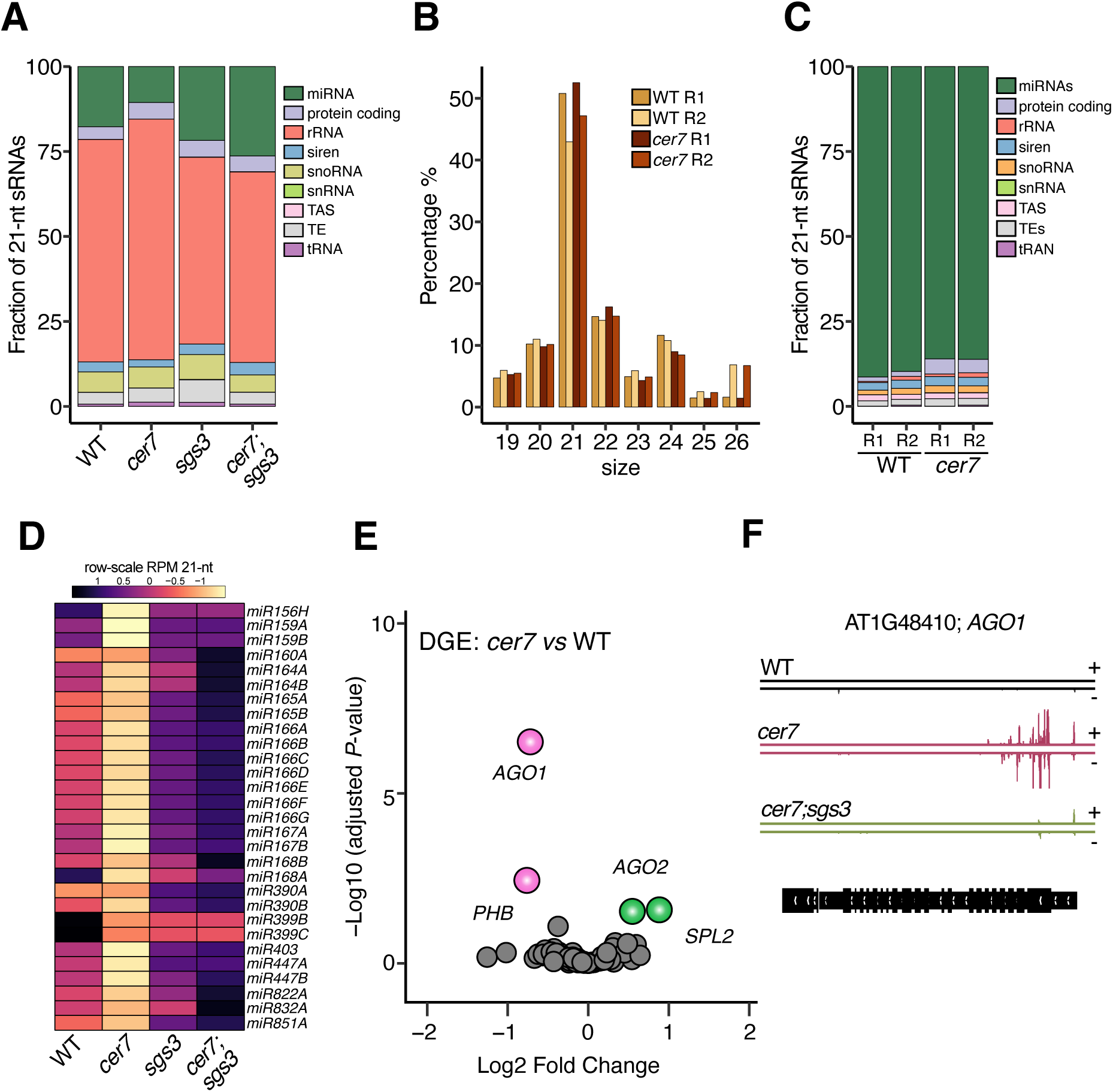
AGO1 loading of a subset of miRNAs is compromised in *cer7*. (**A**) Overview of 21-nt sRNA classes accumulating in 3 DAP seeds of the indicated genotypes. (**B**) Size distribution of sRNAs after AGO1 RIP-seq (**C**) Overview of sRNA classes identified by AGO1 RIP-seq in 2-3 DAP siliques in WT and *cer7*. (**D**) Heat map of miRNAs that overlap with a significantly depleted window of 21-nt sRNAs in AGO1 RIP-seq sRNA-seq *cer7* samples. (**E**) Differential gene expression (DGE) of predicted genes targeted by miRNAs depleted in *cer7*. Pink and green color indicate downregulated and upregulated genes respectively, grey indicate genes with no significant change. (**F**) Genome-browser screen-shot depicting 21-nt siRNAs along *AGO1* gene.

Interestingly, one of the miRNAs significantly depleted from AGO1 is *MIR167*, whose knockout in Arabidopsis was recently shown to cause a maternal sporophytic effect on globular embryo arrest, strikingly similar to *cer7*^14^. This observation suggests that impairment of miRNAs and specifically *MIR167* could explain the globular arrest of *cer7* embryos. To further examine effects on gene expression, we analyzed the transcript abundance of AGO1-depleted miRNA target genes in *cer7* RNA-seq. We found only few miRNA target genes with increased transcript levels in *cer7* (**Fig. 5E**), likely a consequence for the effect being restricted to the embryo, which comprises only a small proportion of the seed. Nevertheless, among the upregulated miRNA target genes we found *SLP2* and *AGO2*, which are regulated by *miR156* and *miR403,* respectively. Both miRNAs were significantly depleted from AGO1 in *cer7 seeds* (**Fig. 5D**). Interestingly, we also observed decreased loading of *miR165/166* and *miR168*, which regulate *PHABULOSA*/*PHAVOLUTA* (*PHB*/*PHV*) and *AGO1*, respectively; however, these miRNA target mRNAs did not increase in abundance, instead, their levels were reduced. A subset of genes targeted by miRNAs are prone to induce secondary siRNAs, including *AGO1* and *PHB*^46,47^. We indeed found enrichment of 21/22-nt siRNAs in *PHB* and *AGO1* in our AGO1 RIP-seq samples of *cer7,* explaining their reduced abundance (**Fig. 5E-F**). These observations substantiate the idea of miRNA depletion through ectopic siRNA competition for AGO1 loading, suggesting that *miR168* and *miR165/166* are likely outcompeted by ectopic 21-nt siRNAs derived from their own target loci.

## Discussion

In this study, we discovered that RNA surveillance in the maternal sporophyte is important to ensure embryo survival, suggesting that the exchange of RNAs between the maternal sporophyte and the embryo is substantially more relevant than what has been generally recognized.

### PTGS-dependent seed abortion in *cer7*

We demonstrated that *cer7* developmental defects are linked to ectopic PTGS, as suppression of PTGS in *sgs3* and *rdr6* mutant backgrounds fully restored seed viability (**Fig. 2A-B**). Despite this suppression, introducing wild-type *SGS3* and *RDR6* alleles through pollen reinstated the seed abortion phenotype, suggesting that PTGS amplification can still occur in the embryo (**Fig. 2A-C**). These findings indicate that a PTGS trigger—likely an aberrant RNA fragment—is generated in the *cer7* sporophyte and subsequently transported to the embryo, where it serves as a template for RDR6-SGS3-mediated siRNA amplification (**Fig. 6**). Validating this idea, expression of *CER7* in the seed coat was sufficient to suppress the seed abortion phenotype of *cer7*, in line with the sporophytic maternal effect in embryo development (**Fig. 3**). Further supporting the role of PTGS in *cer7* embryo lethality, we found that the *cer7;ago1-27* double mutant fully suppressed seed abortion. Analysis of self-pollinated *cer7;ago1-27*/+ heterozygous plants confirmed that AGO1 acts in the embryo to suppress *cer7*-associated embryo arrest (**Fig. 2D-E**). Given that *ago1-27* is a hypomorphic allele deficient in PTGS but not miRNA function^33^, we conclude that maternal PTGS is the primary driver of *cer7* seed abortion.

**Fig. 6:**
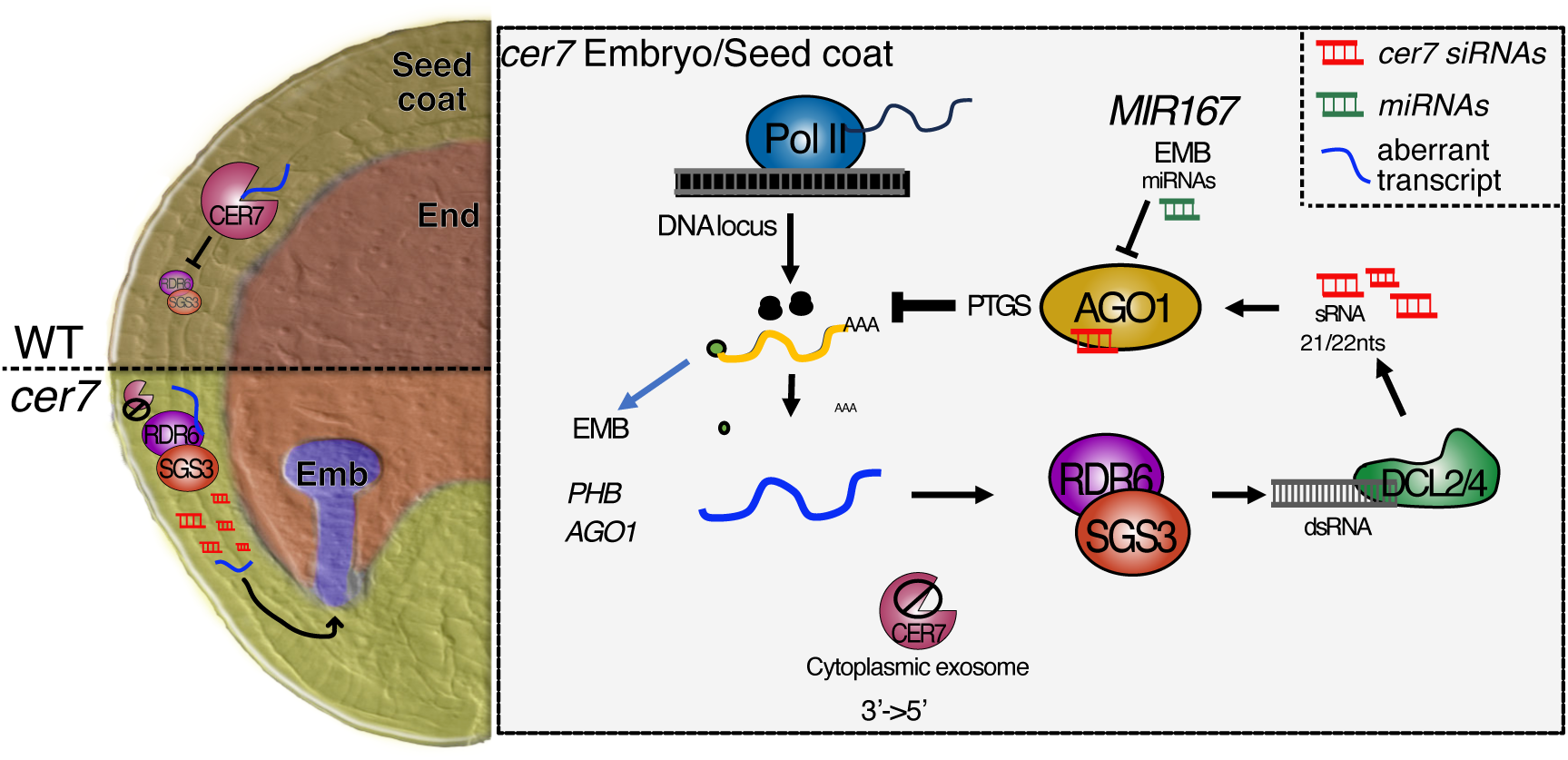
Model of sporophytic-maternally dependent PTGS–mediated embryo arrest in *cer7*. In WT, the exosome clears aberrant transcripts (depicted in blue) preventing their channeling to the PTGS pathway. In *cer7,* aberrant transcripts accumulate in the seed coat and are the templates of RDR6;SGS3 for dsRNA synthesis which are processed by DCL4/2 to produce ectopic 21/22-nt siRNAs (depicted in red). Ectopic siRNAs are loaded into AGO1 and outcompete a subset of miRNAs (depicted in green) for loading onto AGO1, including *MIR167*, which *c*ontributes to the sporophytic maternally controlled embryo arrest in *cer7*. Our data support a model in which an aberrant transcript or subset of transcripts accumulates in *cer7* sporophytic maternal tissues and reaches the embryo, leading to PTGS amplification and seed abortion. Our data also show that PTGS amplification similarly occurs in the sporophytic tissues, likely contributing to an increase in the dosage of ectopic siRNAs reaching the embryo, consistent with the sporophytic maternal effect of *cer7* on embryo development. As a result, genes such as *PHB* and *AGO1* are targeted for ectopic siRNA-mediated silencing, undermining embryonic development.

Nevertheless, RNA-seq and sRNA-seq analyses suggest that mRNA depletion due to PTGS alone does not fully account for the *cer7* phenotype. While only three genes enriched in 21/22-nt siRNAs (*AGO1*, *XTH9*, *BGLU*) exhibited reduced mRNA abundance in *cer7* seeds, AGO1 IP sRNA-seq identified additional targets, including *CER3*, *BGLU37*, and *PHB*. Notably, *PHB* and *AGO1* are expressed in embryonic tissues, and *PHB* mRNA levels were depleted in *cer7* RNA-seq data. Since null *ago1* mutants are inviable but do not exhibit severe early embryo defects^42^, our findings suggest that ectopic siRNA accumulation targeting essential genes, such as *PHB*, contributes to the *cer7* embryo arrest. Weak *ago1* alleles exacerbate embryo defects when *PHB*, *PHV*, and *REV*—all members of the class III HD-Zip transcription factor family—are lost^48^, supporting this hypothesis. Consistently, AGO1 RIP-seq from *cer7* seeds revealed 21-nt secondary siRNAs for *PHB*, *PHV*, and *REV*, as well as ectopic siRNA accumulation for *SAMDC1*, a gene crucial for embryo patterning^49^.

### Defective miRNA loading in AGO1 contributes to *cer7* embryo arrest

Our data furthermore suggest that defective miRNA loading into AGO1 is an additional component of the *cer7* embryo arrest. Supporting this hypothesis, our findings include: 1) a reduction in the proportion of miRNAs in *cer7* sRNA-seq samples from developing seeds; 2) an increase in 21-nt siRNAs from protein-coding genes in *cer7* AGO1 RIP-Seq; 3) significant depletion of key miRNAs crucial for embryo development in *cer7* AGO1 RIP-Seq; 4) *mir167* mutants exhibiting an identical globular embryo arrest with a sporophytic maternal effect^14^; and 5) *dcl1* (*sin1*) mutants, which disrupt miRNA biogenesis, displaying similar maternal-effect embryo arrest^50,51^. These observations suggest that ectopic siRNA accumulation disrupts AGO1 function, leading to impaired miRNA loading and subsequent embryo developmental arrest.

### Maternal sporophyte as a regulator of embryogenesis

The angiosperm seed structure implies a direct maternal influence on embryo development, beyond its mechanical properties^52^. However, the only known symplastic connection between seed coat and zygotic tissues is the suspensor^7^, facilitating auxin transport to the embryo. Given the strong embryo arrest phenotype in *cer7*, along with the absence of observable endosperm defects, our findings support a model in which PTGS precursors and siRNAs move from the maternal seed coat to the embryo via the suspensor, imposing a dosage-sensitive PTGS-dependent arrest (**Fig. 6**). This model is consistent with the observed suppression of *cer7* seed defects upon *CER7* expression in the seed coat. Alternatively, RNA movement could also occur through vesicular trafficking, suggesting that RNA-based maternal control over embryogenesis may involve additional, unexplored transport mechanisms. Determining the nature of these transported RNA molecules and their specific mechanisms of movement represents an exciting avenue for future research.

## Conclusion

This study provides evidence that the RNA exosome plays a crucial role in seed development by preventing the accumulation of aberrant transcripts in maternal seed coat tissues. If left unchecked, these transcripts trigger the production of secondary siRNAs, with deleterious consequences for embryo development. Our findings further indicate that the maternal integuments of the seed function not only as a physical protective structure but also as a permeability barrier where RNA molecules can cross seed compartments, exerting maternal control over embryogenesis. The discovery of CER7 as a key regulator in sporophytic maternal tissues highlights the complex interplay between maternal and embryonic gene regulation, revealing the importance of RNA decay in plant development.

## Methods

### Plant material and growth conditions

Seeds were surface-sterilized using 70% ethanol, followed by three washes with sterile water. They were then sown on ½ MS medium supplemented with 0.5% sucrose and 0.8% agar. After stratification at 4 °C for 2 days, seeds were germinated under long-day conditions (16 h light/8 h dark) at 21 °C. Seedlings were transferred to soil after 10–12 days and grown in phytotrons under long-day conditions (21 °C day/20 °C night, 70% relative humidity, 150 µE light intensity).

All *Arabidopsis thaliana* plants used in this study were in the Columbia-0 (Col-0) genetic background. Mutant alleles were obtained from the Arabidopsis Biological Resource Center (www.arabidopsis.org) and include: *cer7-3* (SAIL_747_B08)^22^, *cer7-4* (GK_089C02)^23^, *rdr6-15* (SAIL_617_H07)^53^, *sgs3-14* (SALK_001394)^54^, *dcl1-14* (SALK_056243) ^36^, and *ago1-27* ^33^.

### Phenotypic analysis of embryo development and DIC microscopy

Homozygous cer7 and wild-type (WT) plants were grown in parallel, emasculated, and pollinated two days later. Siliques were harvested at 2, 3, 5, and 7 days after pollination (DAP), and developing seeds were cleared using a modified Hoyer’s solution consisting of 7.5 g gum Arabic, 100 g chloral hydrate, 5 ml glycerol, and 30 ml water, which was diluted 1:1 with a solution containing 7.5 g gum Arabic, 5 ml glycerol, and 30 ml water. Embryos were visualized using a Leica DM2500 microscope equipped with Nomarski differential interference contrast (DIC) optics.

### Seed abortion and segregation analysis

*cer7* was introgressed into the *rdr6-15*, *sgs3-14*, dcl1-14, and *ago1-27* mutant backgrounds by crossing. Homozygous and heterozygous *cer7* plants and double mutants were identified by PCR genotyping. Crosses were performed by emasculating flowers and pollinating them two days later. Seed abortion was evaluated at 14 DAP using a dissecting microscope, when aborted seeds could be distinguished from WT siblings. At this stage, aborted seeds appeared white, while WT seeds in the same silique were green. All crosses were performed with at least two biological and two technical replicates, using different parent plants and independent experimental batches. Pictures were taken with a Keyence Digital Microscope. Genetic segregation analysis was conducted on heterozygous *cer7* plants identified by PCR genotyping and hand pollinated. Progeny was sown, and individual plants were genotyped by PCR. Segregation ratios were tested against the expected Mendelian ratio for a recessive allele using a Chi-square test. All genotyping primers are listed in **Supplementary Table 4**.

### Feulgen staining

Sample preparation and embedding for Feulgen staining was performed as previously described (Braselton, Wilkinson, and Clulow 1996). Briefly, whole siliques were fixed in ethanol:acetic acid (3:1) overnight, then washed three times in water for 15 minutes each. Samples were incubated in freshly prepared 5 N HCl for 1 hour, followed by three additional 15-minute water washes. Staining was performed in Schiff’s reagent for 3 hours, followed by three washes in cold water and sequential 10-minute washes in increasing concentrations of ethanol (10%, 30%, 50%, 70%, 90%). The samples were then incubated in 100% ethanol overnight. Embedding was performed using a graded ethanol:LR White resin series (3:1, 1:1, 1:3) for 1 hour each, followed by overnight incubation in pure LR White. Samples were mounted in LR White with accelerator and polymerized overnight at 60°C. Samples were imaged using a LEICA Stellaris 8 DIVE microscope in multiphoton mode, with excitation at 800 nm and detection in the 563– 668 nm emission range.

### Construction of CER7-GFP transgenic plants

Regions upstream of the translation start sites of *dVPE* (-2,976 bp; AT3G20210), *BAN* (-191 bp), and *RPS5A* (-1,660bp; AT3G11940) were cloned into the pB7FWG.2 vector ^55^, replacing the CaMV 35S promoter. The pB7FWG.2 vector was linearized with *Sac*I and *Spe*I, and the upstream fragments were inserted using the Takara In-Fusion Snap Assembly Kit (TAKARA, 638945) to generate new destination vectors. The *CER7* coding sequence (1317 bp) was amplified from cDNA and recombined into pDONR221 using the Gateway BP Clonase II Enzyme Mix (ThemoFisher, 11789100). Final constructs, *pdVPE::CER7-GFP*, *pBAN::CER7-GFP*, and *RPS5A::CER7-GFP*, were obtained via LR recombination (ThermoFisher, 11791020) between CER7_pDONR221 and the corresponding promoter-containing destination vectors. See **Supplementary Table 4** for the list of promoters used for cloning.

Heterozygous *cer7* plants were transformed with the final constructs using the floral dip method ^56^. Transgenic plants were selected based on Basta resistance and PCR. GFP fluorescence was assessed using a Leica DM2500 fluorescence microscope. Transgenic heterozygous and homozygous *cer7* lines were identified by PCR genotyping. Complementation analysis was carried out in homozygous *cer7* backgrounds using at least seven independent transgenic lines.

### Confocal microscopy

Imaging of *pdVPE::CER7-GFP, pBAN::CER7-GFP*, and *RPS5A::CER7-GFP* expression was done using a LEICA Stellaris 8 Dive microscope with an excitation of 488 nm and an emission range of 493–551 nm. Developing seeds were mounted in 10% glycerol and fluorescence was observed.

### RNA-seq

Wild-type (WT), *cer7-4*, *sgs3-14*, and *cer7-4;sgs3-14* double mutant plants were manually self-pollinated. Siliques were dissected at 3 days after pollination (DAP), and approximately 500 seeds per genotype were collected and stored in RNAlater solution (ThermoFisher, AM7021). Total RNA was extracted using TRIzol Reagent (Invitrogen, 15596018) according to the manufacturer’s instructions. RNA samples were treated with DNase I at 37 °C for 1 hour (ThermoFisher, EN0521), followed by heat inactivation at 65 °C for 10 minutes and DNase removal by TRIzol extraction. mRNA libraries were prepared using the NEBNext Ultra II DNA Library Prep Kit (NEB, E7645S) in combination with the NEBNext Poly(A) mRNA Magnetic Isolation Module (NEB, E7490S). Sequencing was performed on a NextSeq 1000 platform using 150-bp paired-end (PE) mode.

PE reads were trimmed using Trim Galore by removing 5 bp from the 5′ end and 20 bp from the 3′ end. Trimmed reads were aligned to the *Arabidopsis thaliana* reference genome (TAIR10) using HISAT2 ^57^. Mapped reads were sorted and indexed with SAMtools ^58^. Gene-level read counts were obtained using HTSeq-count, and expression values were normalized to transcripts per million (TPM) using StringTie. Differential gene expression analysis was performed with DESeq2 (v1.42.0) ^59^ in R (v4.3.0). Genes with an absolute log2 fold change (LFC) ≥ 0.8 and a false discovery rate (FDR) < 0.05 were considered significantly differentially expressed. Data visualization was carried out using the ggplot2 package in R.

### sRNA-seq

sRNA libraries were generated for wild-type (WT), *cer7-4*, *sgs3-14*, and *cer7-4;sgs3-14* using same DNAse-treated RNA used for mRNA-seq libraries referred above. 1-2ug of DNAse-treated RNA was loaded into a denaturing acrylamide gel at 16% and gels bands between 16- and 30-nt were excised and small RNA enriched samples were recovered using DNA gel elution buffer from NEB and a Cellulose Acetate Membrane column. Libraries were made using the NEBNext Small RNA Library Prep Set for Illumina (New England Biolabs) following the manufacturer’s instructions. Sequencing was performed on a NovaSeq 6000 platform using 150-bp paired-end (PE) mode. Only one read of the pair was used in further analyses.

Adapters were removed from the first read of each 150-bp PE library using Cutadapt, and reads between 19–26 nt in length were retained for further analysis. Processed reads were aligned to the *Arabidopsis thaliana* reference genome (TAIR10), and small RNA loci were annotated using ShortStack ^60,61^ with the parameters --mincov 2 and --mmap f. To identify regions with differential small RNA accumulation, 21-nt and 22-nt reads were extracted from BAM files using SAMtools ^58^, and coverage across 100-bp genomic windows was calculated using the bedtools multicov function. Differential accumulation of small RNAs across windows was assessed with DESeq2 ^59^ in R (v4.3.0). Windows with an absolute log2 fold change (LFC) ≥ 0 and a false discovery rate (FDR) < 0.05 were considered significant and merged if they were within 100 bp of each other using the bedtools merge function. Significant windows were annotated to TAIR10 genome features using bedtools intersect, and visually inspected in a genome browser to confirm genotype-specific enrichment or depletion. Data visualization was performed using the ggplot2 package in R.

### AGO1 RIP-seq

Siliques closest to the inflorescences of homozygous *cer7* and wild-type (WT) plants were removed, and newly formed siliques were harvested five days later, corresponding to an estimated 2–3 days after self-pollination. For each replicate, 500 mg of siliques were collected and flash-frozen in liquid nitrogen. Endogenous AGO1 immunoprecipitation was performed as previously described ^62^. Briefly, tissue was ground to a fine powder using a mortar and pestle, resuspended in three volumes of crude extract buffer (50 mM Tris-HCl, pH 7.5; 150 mM NaCl; 10% glycerol; 5 mM MgCl₂; 0.1% IGEPAL; 5 mM DTT; and 1× cOmplete™ Protease Inhibitor Cocktail), and incubated at 8 rpm for 20 minutes at 4 °C. The crude extract was centrifuged at 20,000 × g for 15 minutes, repeated 2–3 times until clarified.

Equal amounts of clarified extract were incubated with AGO1 antibody (Agrisera, AS09527) pre-bound to PureProteome Protein A magnetic beads (Millipore, LSKMAGA02) for 3 hours at 4 °C on a rotating carousel (10–12 rpm). Immune complexes were washed four times with crude extract buffer, and AGO1-associated RNA was extracted directly from the beads using TRIzol Reagent (Invitrogen, 15596018) following the manufacturer’s protocol. Small RNA library preparation and downstream analysis were performed as described above, omitting the initial small RNA enrichment step. Sequencing was performed on a NextSeq 1000 platform.

## Data availability

Sequencing data generated in this study are available in the Gene Expression Omnibus in NCBI under the accession numbers GSE295374 for RNA-seq and GSE295375 for sRNA-seq.

## Acknowledgments

We thank Daniela Barro-Trastoy and Kai Wang for helpful comments on the manuscript.

## Author contributions

G.D.T.D.L. and C.K. conceptualized the project, developed the methodology and provided supervision. G.D.T.D.L., M.S.T., V.V., and U.K. conducted experiments, G.D.T.D.L. performed bioinformatic analyses. C.K. acquired funding and administered the project. G.D.T.D.L. and C.K. wrote the original paper draft. All authors reviewed and edited the paper.

