## Supplemental Figures for "Maternal control of RNA decay safeguards embryo development"

**The PDF file includes:**

Tables S1 to S3

Figs. S1 to S9

**Other Supplementary Materials for this manuscript include the following:**

Data S1 to S3

**Supplementary Tables**

| **Supplementary Table 1.** Summary RNA-seq data. Developing seeds 3 DAP | | | | | | | | |
|---|---|---|---|---|---|---|---|---|
| | | | | **hisat2 alignment** | | | | |
| **genotype** | **sample replicate** | **color code** | **Total unique Raw reads (R1+R2)** | **Total unique Raw reads (R1+R2)** | **Aligned 0 times** | **Aligned 1 time** | **Aligned >1 times** | **Total alignment rate** |
| *cer7-4/-* | cer7 R1 | | 42.797.938 | 21.277.234 | 525248 (2.47%) | 17555254 (82.51%) | 3196732 (15.02%) | 98,48% |
| | cer7 R2 | | 40.309.854 | 20.091.742 | 470908 (2.34%) | 17212687 (85.67%) | 2408147 (11.99%) | 98,61% |
| | cer7 R3 | | 83.895.834 | 41.609.263 | 1116394 (2.68%) | 35828720 (86.11%) | 4664149 (11.21%) | 98,32% |
| *sgs3-14/-* | sgs3 R1 | | 96.704.242 | 47.981.077 | 1369175 (2.85%) | 39512910 (82.35%) | 7098992 (14.80%) | 98,14% |
| | sgs3 R2 | | 97.180.336 | 48.333.652 | 1235413 (2.56%) | 33293303 (68.88%) | 13804936 (28.56%) | 98,45% |
| | sgs3 R3 | | 95.767.222 | 47.616.277 | 1244472 (2.61%) | 42984338 (90.27%) | 3387467 (7.11%) | 98,37% |
| *sgs3-14/-; cer7-4/-* | sgs3cer7 R1 | | 43.933.824 | 21.917.616 | 549588 (2.51%) | 19617121 (89.50%) | 1750907 (7.99%) | 98,46% |
| | sgs3cer7 R2 | | 136.557.724 | 68.073.762 | 1793208 (2.63%) | 46628770 (68.50%) | 19651784 (28.87%) | 98,40% |
| | sgs3cer7 R3 | | 122.046.250 | 60.822.450 | 1499010 (2.46%) | 41200582 (67.74%) | 18122858 (29.80%) | 98,56% |
| Col WT | WT R1 | | 73.941.874 | 36.796.134 | 968967 (2.63%) | 33087085 (89.92%) | 2740082 (7.45%) | 98,35% |
| | WT R2 | | 49.317.868 | 24.590.744 | 548920 (2.23%) | 22380067 (91.01%) | 1661757 (6.76%) | 98,63% |
| | WT R3 | | 46.085.514 | 22.965.100 | 535636 (2.33%) | 17452063 (75.99%) | 4977401 (21.67%) | 98,64% |

| **Supplementary Table 2.** Summary sRNA-seq data. Developing seeds 3 DAP | | | | | | | |
|---|---|---|---|---|---|---|---|
| **genotype** | **sample replicate** | **color code** | **18-30nt** | | **Trimmed raw reads 19-16nt** | **Total mapped 19-26nt** | **Mapping %** |
| | | | **Total reads processed R1** | **Reads passing trimming filters R1** | | | |
| *cer7-4/-* | cer7 R1 | | 22.084.951 | 18.751.841 | 14.977.781,00 | 13.754.657 | 91,83% |
| | cer7 R2 | | 17.650.699 | 15.598.131 | 12.846.321,00 | 12.096.366 | 94,16% |
| | cer7 R3 | | 31.218.807 | 27.391.195 | 20.548.030,00 | 19.502.645 | 94,91% |
| *sgs3-14/-* | sgs3 R1 | | 40.603.526 | 34.974.211 | 30.970.165,00 | 29.031.562 | 93,74% |
| | sgs3 R2 | | 21.750.633 | 18.921.084 | 15.644.498,00 | 14.843.925 | 94,88% |
| | sgs3 R3 | | 27.175.358 | 24.012.911 | 19.915.276,00 | 18.591.402 | 93,35% |
| *sgs3-14/-; cer7-4/-* | sgs3cer7 R1 | | 31.878.308 | 28.007.754 | 25.419.772,00 | 23.740.355 | 93,39% |
| | sgs3cer7 R2 | | 29.383.397 | 26.632.177 | 23.313.734,00 | 21.843.371 | 93,69% |
| | sgs3cer7 R3 | | 17.898.486 | 16.505.193 | 14.477.458,00 | 13.683.859 | 94,51% |
| Col WT | WT R1 | | 34.604.248 | 32.530.105 | 26.395.400,00 | 25.210.860 | 95,5% |
| | WT R2 | | 39.133.930 | 36.202.909 | 30.133.762,00 | 28.851.901 | 95,70% |
| | WT R3 | | 43.898.777 | 37.481.536 | 33.373.569,00 | 30.960.280 | 92,76% |

| **Supplementary Table 3.** Summary AGO1 RIP-seq data. Siliques 3 DAP | | | | | | | | |
|---|---|---|---|---|---|---|---|---|
| **Sample** | **replicate** | **Color code** | **Trimming summary (18--30nt)** | | **Mapping** | | | |
| | | | **Total R2 processed** | **passing filters** | **19-26nt reads** | **Mapped** | **Unmapped** | **Mapping %** |
| *cer7-1* | cer7 R1 | | 43,352,066 | 31,371,831 (72.4%) | 29.241.256 | 26.723.117 | 2.518.139 | 91,39 |
| | cer7 R2 | | 70,324,366 | 34,035,494 (48.4%) | 30.337.927 | 27.406.777 | 2.931.150 | 90,34 |
| WT | WT R1 | | 60,045,705 | 42,150,778 (70.2%) | 39.348.821 | 36.507.360 | 2.841.461 | 92,78 |
| | WT R2 | | 72,168,116 | 27,880,827 (38.6%) | 24.812.728 | 22.677.707 | 2.135.021 | 91,40 |

**Supplementary Figures**

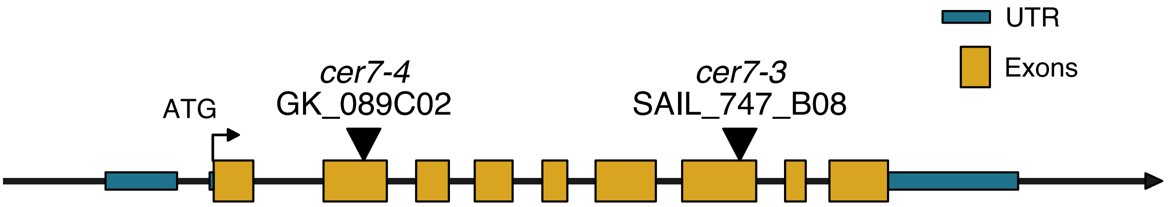


**Supplementary Fig. 1: Schematic representation of the *RRP45B/CER7* gene (AT3G60500) and the mutant alleles used in this study.** Yellow boxes indicate exons, blue boxes indicate untranslated regions (UTRs), and black lines indicate introns and intergenic regions. Black triangles represent estimated regions of T-DNA insertions. Both T-DNA insertions are reported as loss-of-function alleles (Hooker et al. 2007; Lange et al. 2019).

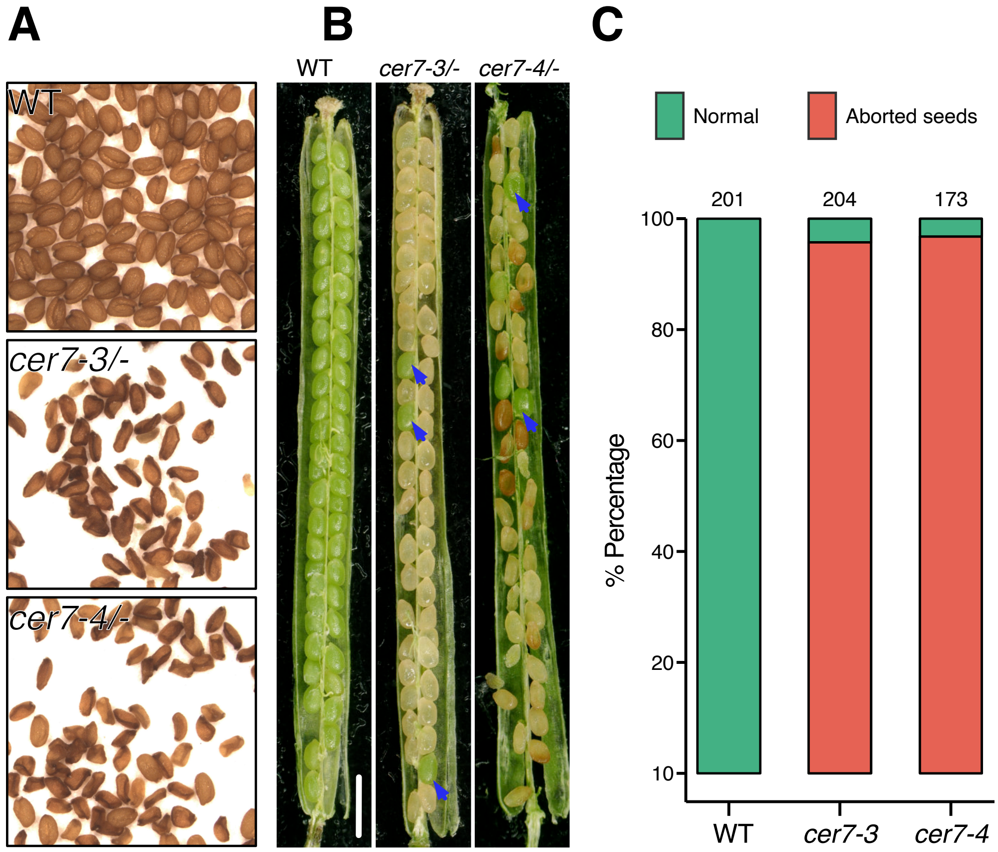


**Supplementary Fig. 2: Seed abortion phenotype in *cer7* alleles.** (**A**) Mature seed abortion phenotypes. (**B**) Siliques at 14 DAP. Bar=1000µm. (**C**) Quantification of seed abortion phenotypes from B. Blue arrow indicates WT sibling of the same silique.

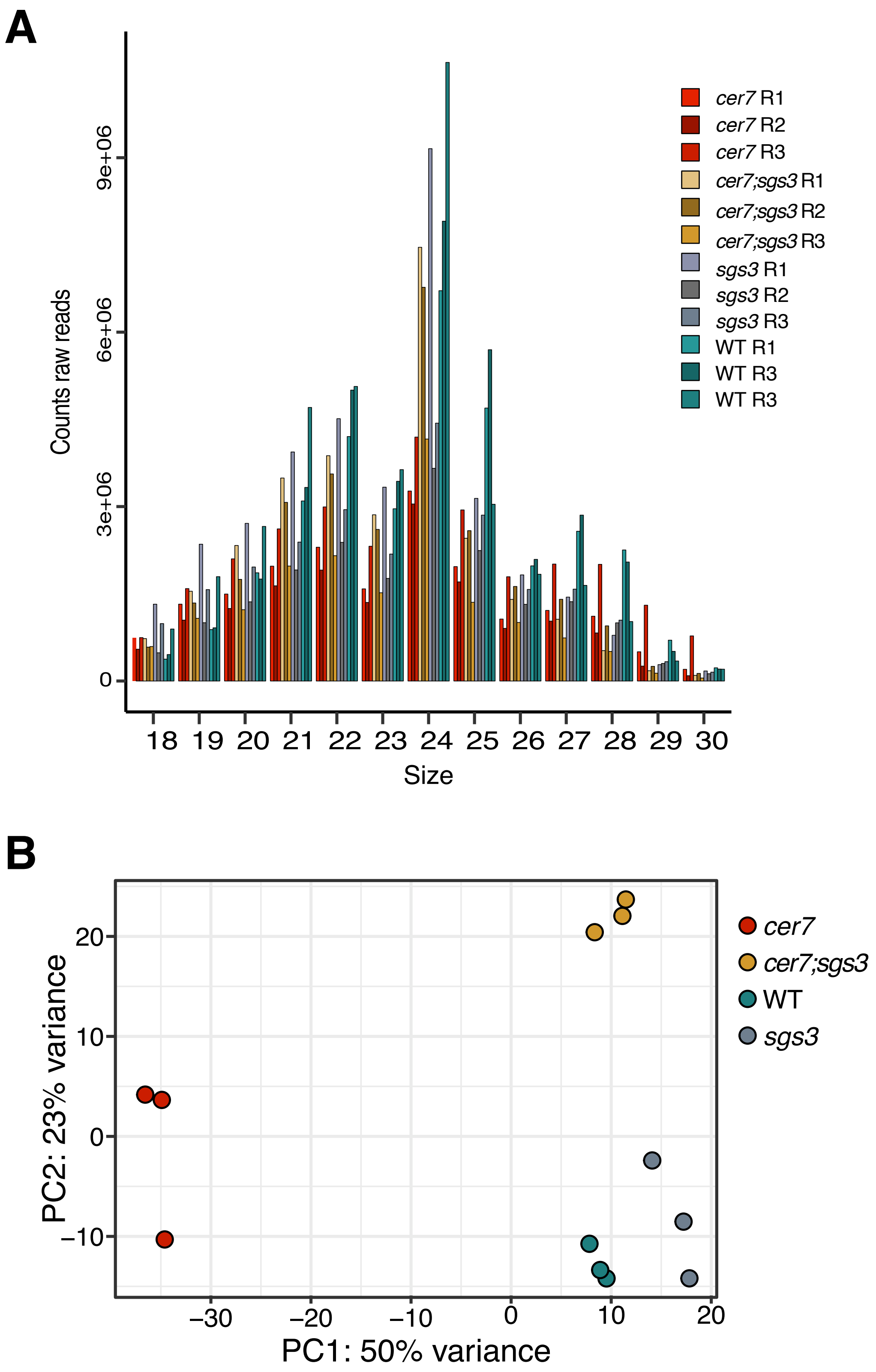


**Supplementary Fig. 3: Small RNA-seq of 3 DAP seeds.** (**A**) Small RNA size distribution of raw reads. (**B**) Principal component analysis (PCA) of the samples based on mapped reads of 21/22-nt size.

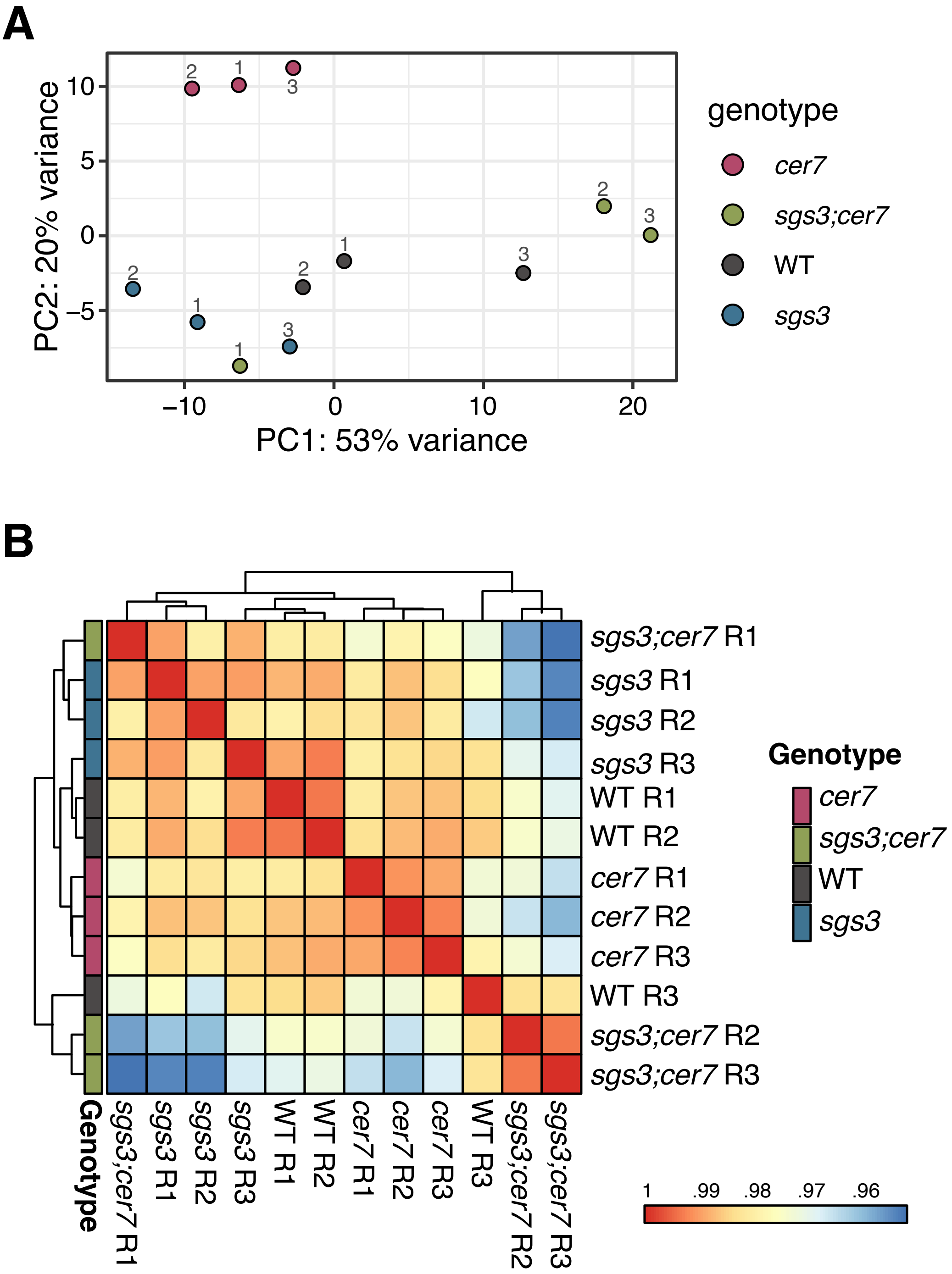


**Supplementary Fig. 4: RNA-seq of 3 DAP seeds.** (**A**) Principal component analysis (PCA) of RNA seq samples. (**B**) Correlation matrix heatmap of transcript expression across samples. PCA, cluster dendrogram, and Spearman correlation coefficient heatmap are based on normalized TPM (transcripts per million mapped reads) values of expressed transcripts.

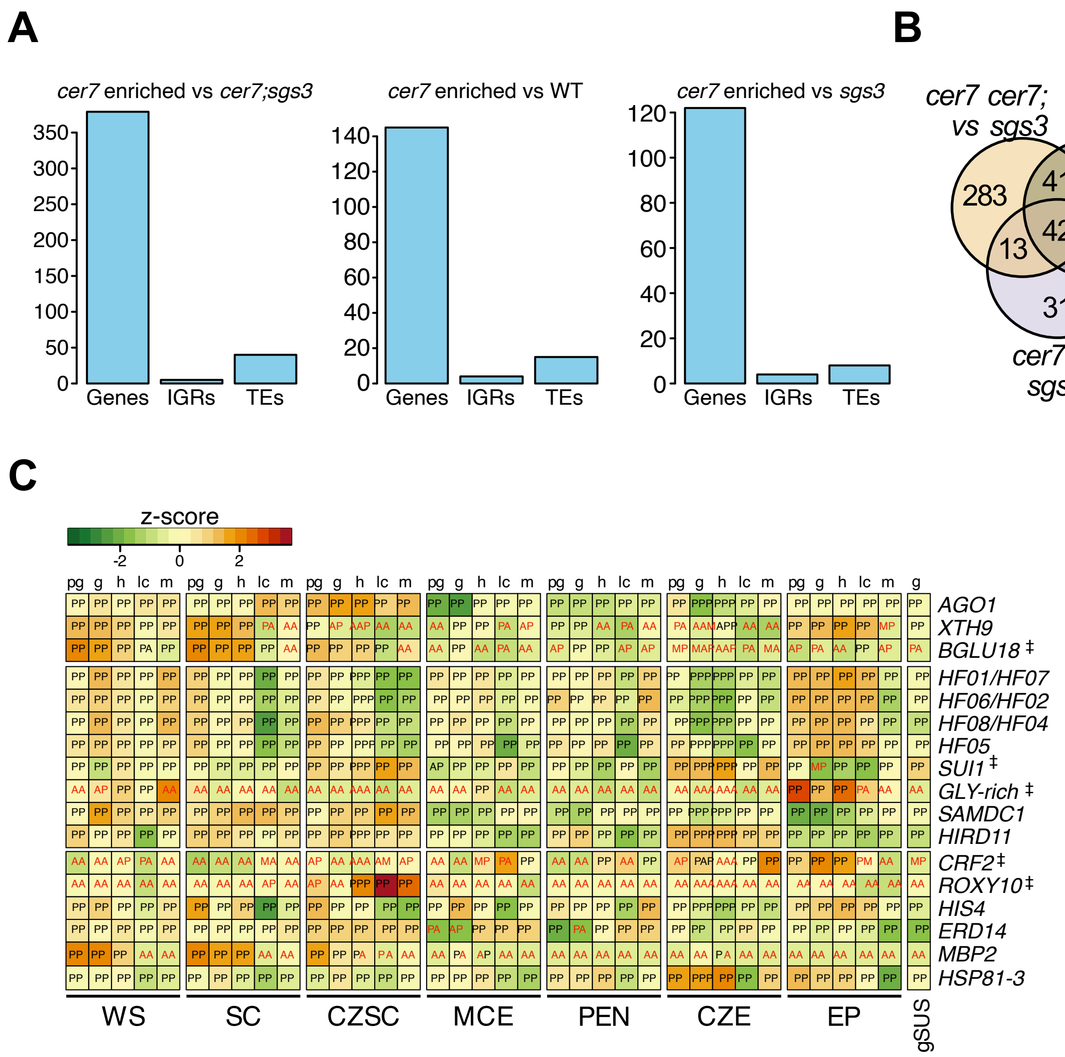


**Supplementary Fig. 5: Identification of 21/22-nt enriched regions in *cer7*.** (**A**) Annotation of significantly enriched 21/22-nt sRNA windows in *cer7* in the indicated comparisons. (**B**) Overlap of genes with enriched 21/22-nt sRNA windows identified in (A). (**C**) Expression of genes enriched for 21/22-nt sRNAs in *cer7* across different seed compartments at different stages of development. LFC, log fold change, FDR, false discovery rate. Meanbase from DGE analysis. ‡ Genes with low or no expression in the embryo. Letters within heatmap boxes indicate MAS5 detection calls: P, present; A, absent; M, marginal. Letters above heatmap boxes indicate developmental stages: pg, preglobular; g, globular; h, heart-stage; lc, linear cotyledon; m, mature green. WS, whole seed; SC, seed coat; CZSC, chalazal seed coat; MCE, micropylar endosperm; PEN, peripheral endosperm; CZE, chalazal endosperm; EP, embryo proper; SUS, suspensor (Belmonte et al., 2013).

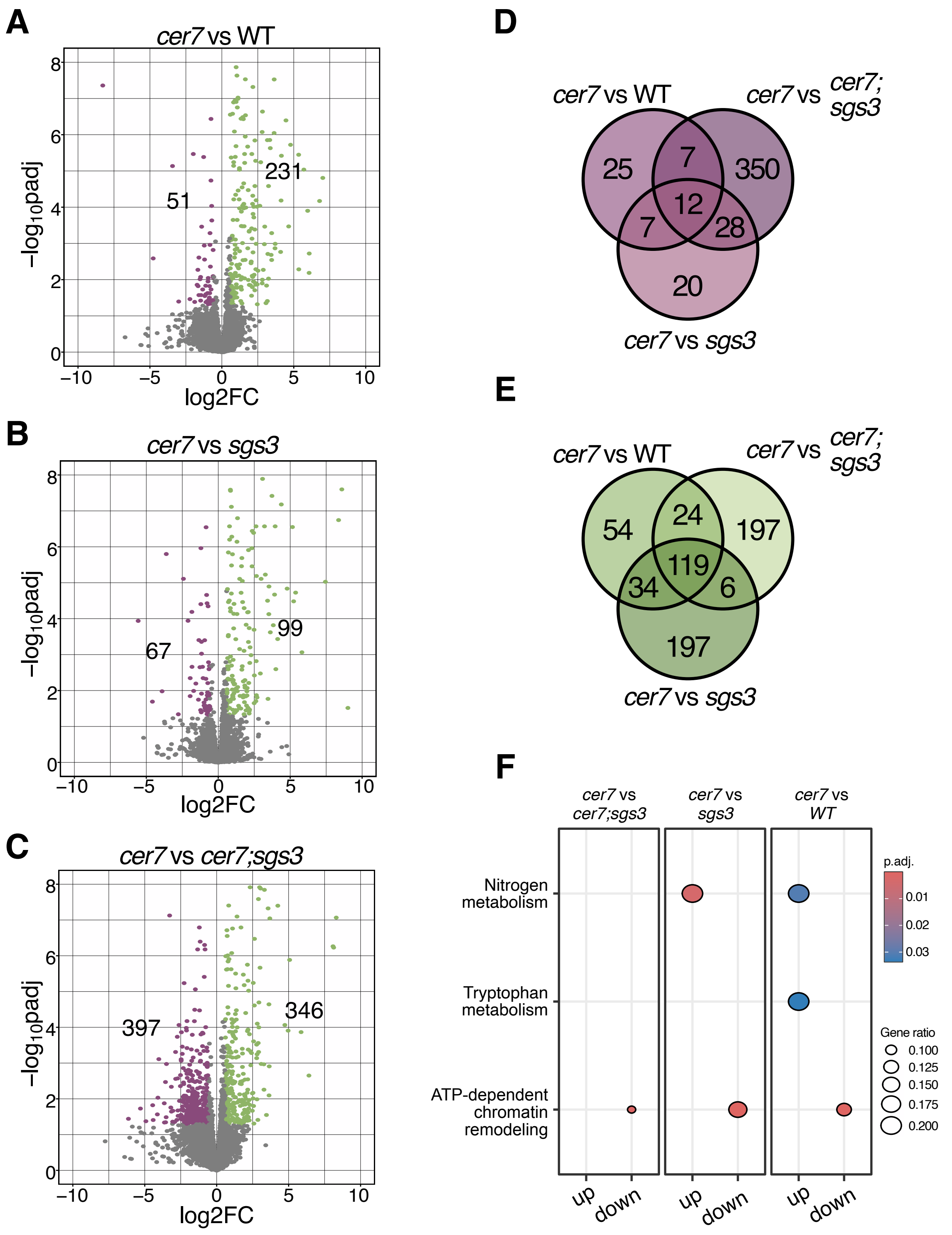


**Supplementary Fig. 6: Differential gene expression (DGE) analysis in 3 DAP seeds.** (**A-C**) Up- and Downregulated genes (LFC ±0.58; FDR <0.05) in the indicated genotype comparisons. (**D**) Overlap of downregulated genes in *cer7*. (**E**) Overlap of upregulated genes in *cer7*. (**F**) Gene ontology analysis by molecular function of deregulated genes.

**A**

**RNA-seq 3DAP developing seeds**

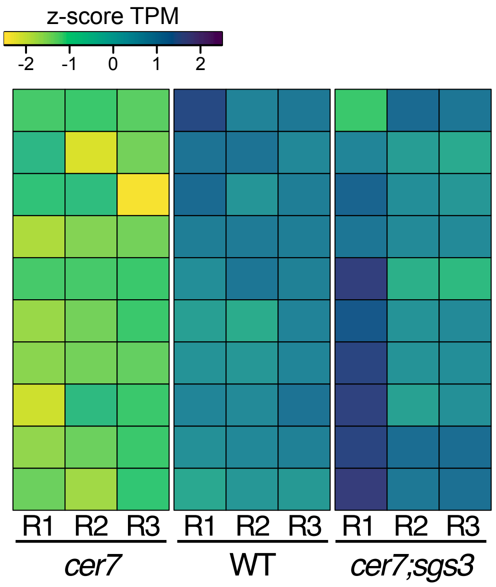


Differentially expressed genes

| | MeanBase | *cer7* vs WT | | *cer7* vs *cer7;sgs3* | |
|---|---|---|---|---|---|
| | | LFC | FDR | LFC | FDR |
| *SBT4.13*‡ | 153,5 | -1,5 | <0.01 | -1,2 | <0.05 |
| *BGLU18*‡ | 102962,9 | -3,4 | <0.01 | -2,3 | <0.01 |
| *BGAL3* | 8089,2 | -1,4 | <0.01 | -1,4 | <0.01 |
| *PFD* | 1821,9 | -3,2 | <0.01 | -3,0 | <0.01 |
| *NITR2*‡ | 156,2 | -1,7 | <0.05 | -1,5 | <0.05 |
| *PHB* | 181,7 | -0,8 | <0.01 | -1,1 | <0.01 |
| *AGO1* | 6174,7 | -0,8 | <0.01 | -1,0 | <0.01 |
| *cis-NAT*‡ | 113,8 | -0,8 | <0.05 | -0,8 | <0.01 |
| *NIT2*‡ | 598,9 | -1,1 | <0.01 | -1,6 | <0.01 |
| *XTH9* | 10655,4 | -0,7 | <0.01 | -1,4 | <0.01 |

**B**

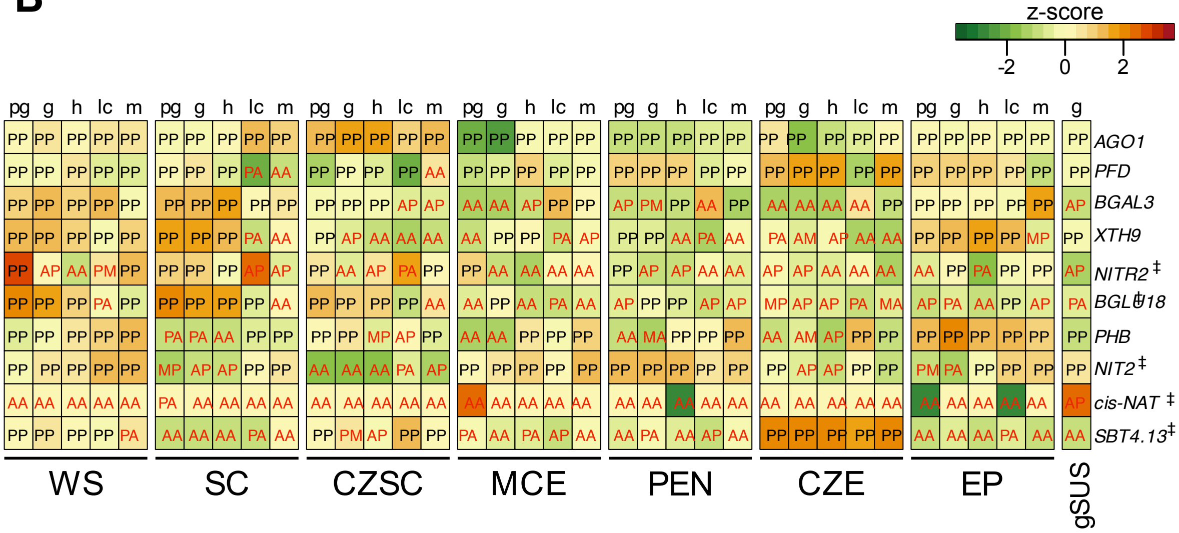


**Supplementary Fig. 7: Downregulated genes in *cer7* seeds at 3 DAP.** (**A**) RNA-seq heat map of downregulated genes in *cer7* at 3 DAP. (**B**) Expression of downregulated genes in different seed compartments at different stages of development. LFC, log fold change, FDR, false discovery rate. Meanbase from DGE analysis. ‡ Genes with low or no expression in the embryo. Letters within heatmap boxes indicate MAS5 detection calls: P, present; A, absent; M, marginal. Letters above heatmap boxes indicate developmental stages: pg, preglobular; g, globular; h, heart-stage; lc, linear cotyledon; m, mature green. WS, whole seed; SC, seed coat; CZSC, chalazal seed coat; MCE, micropylar endosperm; PEN, peripheral endosperm; CZE, chalazal endosperm; EP, embryo proper; SUS, suspensor (Belmonte et al., 2013).

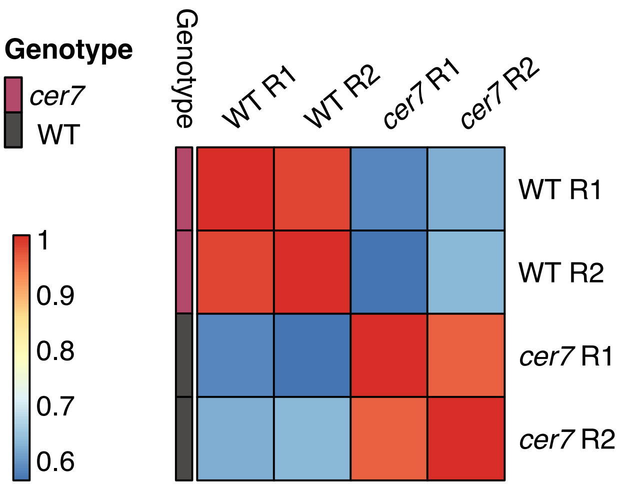


**Supplementary Fig. 8: Small RNA-seq following AGO1-IP of 2-3 DAP siliques.** Spearman correlation heatmap based on VST-transformed counts (21-22nt reads).

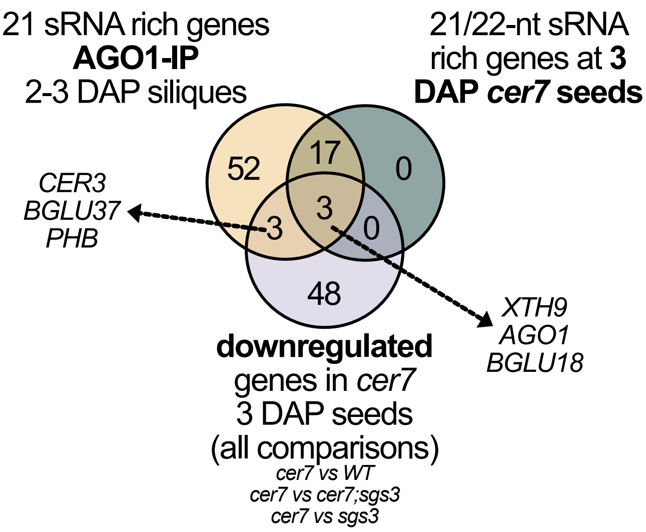


**Supplementary Fig. 9: Overlap of downregulated genes and 21/22-nt enriched genes in *cer7*.** Genes enriched for 21/22-nt sRNAs were determined based on sRNA-seq data of developing seeds at 3 DAP and AGO1-IP sRNA-seq data from 2-3 DAP siliques.
